# Evolution of passwords for cost-free honest signalling between symbionts and hosts

**DOI:** 10.1101/065755

**Authors:** Marco Archetti, J. Benjamin Miller, Douglas W. Yu

## Abstract

Honest communication between potential partners with conflicting interests is generally thought to require costly signals. Costly signalling can explain partner choice when it is possible to link a strategic cost to an individual’s quality, like in mate choice. However, in mutualisms, it is usually impossible to link a cost to the likelihood that a potential partner will behave cooperatively in the future. In fact, signals like Nod factors in rhizobial bacteria, which form symbioses with leguminous plants, are evidence of cost-free, honest signals in situations of potential conflict. How can such a signalling system evolve? We use a population-genetics model to show that a cost-free, honest signal can evolve when the receiver is under *soft selection*, which is when high juvenile mortality does not lead to a corresponding reduction in fitness, a common occurrence in many species. Under soft selection, senders evolve increasingly complex messages of identity, a system akin to a password or a lock and key. Thus, a symbiont can signal that it shares a coevolutionary history with a potential host, and if that history is mutualistic, then the host can believe that the symbiont is mutualistic. Password signalling might also explain the acquisition of some defensive symbionts and the evolution of complex species-recognition signals in mate choice.

> “*…Say now Shibboleth: and he said Sibboleth: for he could not frame to pronounce it right. Then they took him, and slew him…*.” — Judges 12:6, King James Version

## Introduction

### A ‘mechanism-design’ problem in mutualisms

Let us think of communication between symbionts and hosts as a signalling problem. While both mutualistic and parasitic partners have an incentive to enter a host, parasites decrease a host’s fitness. Hence, interests are not aligned, similar to what is found in mate choice, where males of both high and low quality have an incentive to mate, and females have an incentive to choose high-quality males. The host faces a ‘mechanism-design’ problem: how to design a signalling system in which a mutualistic symbiont can uniquely identify itself as a mutualist to a host. In other words, a symbiont must be able to signal its ‘mutualistic identity’ (i.e. that the signaller will be reliably ‘nice’ to the host).

The difficulty with using the classic mechanism of costly signals (Grafen 1990; Maynard Smith and Harper 2003) is that it is not readily apparent how a mutualistic nature can be correlated with the strategic cost that is required for the maintenance of honest signalling in situations of non-aligned interests. Costly signalling is arguably possible in a few cases like big, symmetrical flowers, in which a signal of vigour *per se* can honestly signal that the flower is likely carrying high amounts of rewards. But in most mutualisms, the mere demonstration that a symbiont is *vigorous* does not demonstrate that the symbiont will also *behave mutualistically* in the future (Edwards and Yu 2007). Thus, a strategic cost does not seem to provide a good explanation for honest signalling in mutualisms.

Nonetheless, evolution is exceedingly clever and seems to have solved this mechanism-design problem for Nod factors in legume-rhizobia symbioses (Oldroyd 2013). When the root of a leguminous plant perceives Nod factor from rhizobial bacteria, the root initiates a signalling cascade that results in the formation of an ‘infection thread’ to allow the bacteria to colonize the host plant. All sorts of bacteria would benefit from gaining entry to a root, so there is a strong temptation for pathogens to counterfeit Nod factor. Nonetheless, only rhizobial bacteria appear to make Nod factors that are successfully recognised by the plant to initiate signalling and infection events.

There are three general theoretical classes of honest signalling. We can rule out the ‘costly, honest’ signal explanation of Nod factor, since there is no obvious *strategic* cost (additional to the mere cost of producing the necessary molecules [Maynard Smith and Harper, 2003]) to Nod factor. That is, Nod factor is not a bundle of ammonium molecules, serving as evidence that the bacterium is capable of fixing nitrogen. We can also exclude ‘cheap talk’ signalling (Crawford and Sobel 1982), which requires shared (partially aligned) interests between signaller and receiver individuals, such as occurs between kin. But in horizontally transmitted mutualisms, host and symbiont are different species and disperse separately, erasing shared interests. The third class, ‘verifiable information,’ requires that the signal be true on its face. In biology, verifiable-information signals exist within the concept of the *index* (Maynard Smith and Harper 2003). For instance, claw marks high up on a tree trunk are a self-evident, and thus believable, signal of a tiger’s large size. For Nod factor to be an index, it would need to be a unique by-product of the same biochemical pathway that leads to the quality being sought (nitrogen fixation), so that the mere presence of Nod factor would indicate a mutualistic symbiont, but this is not the case. Nod factor can, in principle, be synthesized by non-nitrogen-fixing rhizobial bacteria, which would appear to rule out verifiable-information signals as well, but we will show how Nod factor can be included in this class.

### Cost-free, honest signals: passwords and Nod factors

To start, we propose that Nod factors be thought of as ‘password signals,’ cost-free messages of arbitrarily high complexity that can honestly convey identity. The idea that Nod factors serve as passwords arises naturally from the many descriptions of Nod factors as chitin-based chains adorned with multiple ‘decorations’ that vary across rhizobial species (**Figure 1**) and the portrayal of Nod factors and Nod-factor receptors acting in a ‘lock-and-key’ manner (Parniske and Downie 2003), such that different Nod factors are accepted by different plant species (Perret et al. 2000), leading to a high degree of species-specificity in rhizobia-hostplant associations.

**Fig. 1.**
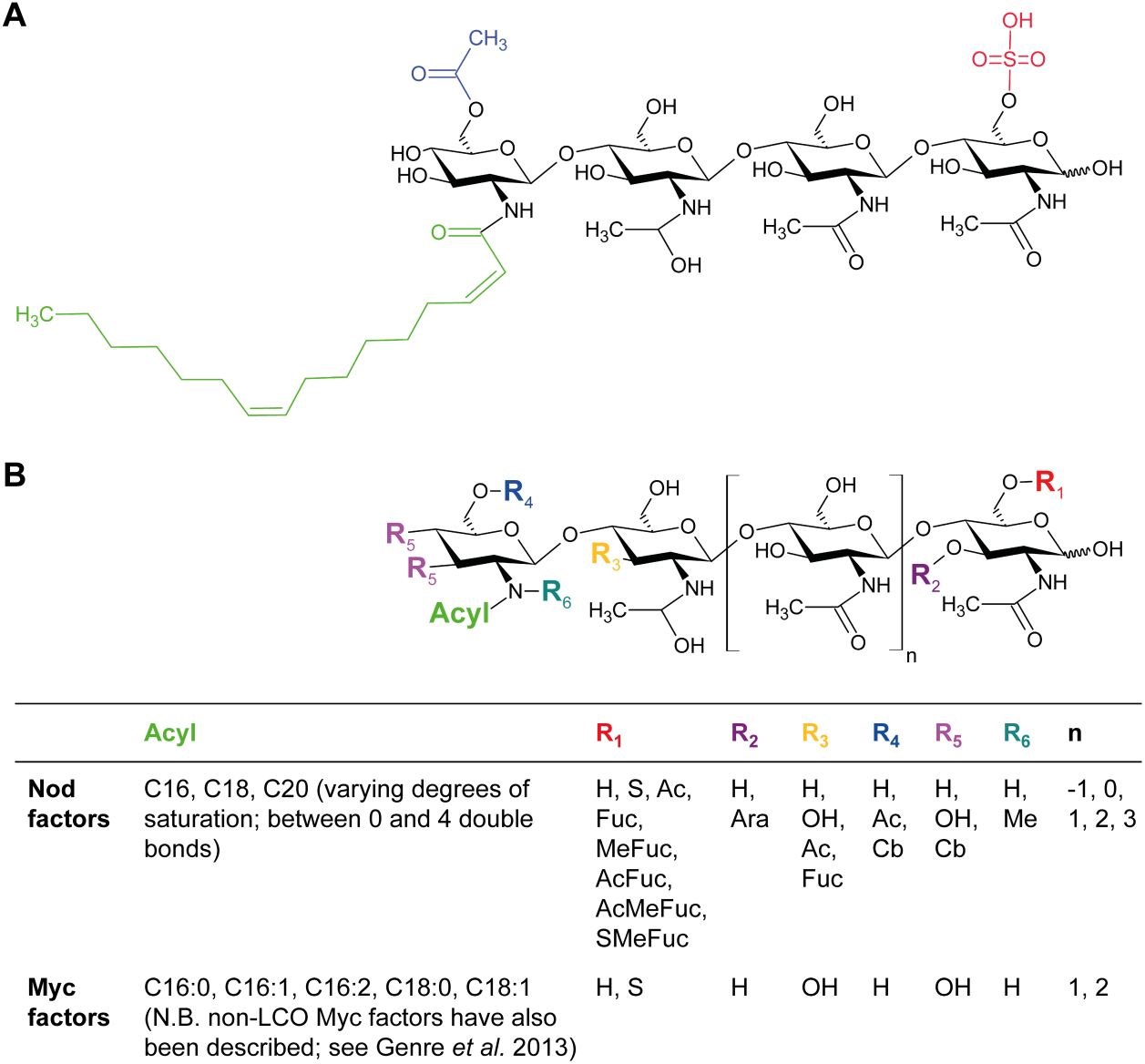
Structures of Nod and Myc factors. **A**. The major Nod factor produced by *Sinorhizobium meliloti* consists of four N-acetylglucosamine residues (black) and a C16:2 acyl group (green). This lipochitooligosaccharide (LCO) backbone is further decorated with sulphate (red) and acetyl (blue) groups. **B**. Generalised structure of Nod and Myc factors. The table shows some of the major decorations and length variants that have been characterised to date. **Ac:** acetyl, **Ara:** arabinosyl, **Cb:** carbamoyl, **Fuc:** fucosyl, **H:** hydrogen, **Me:** methyl, **OH:** hydroxyl, **S** sulphate, **AcFuc:** acetylated fucose, **MeFuc:** methylfucose, **AcMeFuc:** acetylated methylfucose, **SmeFuc:** sulphated methylfucose. Figure adapted from Perret et al. (2000), Wais et al. (2002), and Miller and Oldroyd (2011).

Nod factors are famously variable in structure, with a lipochitooligosaccharide (LCO) backbone of three to five N-acetylglucosamine residues to which multiple different “decorations” can be added, e.g. addition of an acetyl, methyl, sulphate, or sugar moiety (**Figure 1**).

The set of Nod factor decorations varies across rhizobial species and biovars, as does the length of the LCO backbone and the acyl (fatty-acid) chain added to the non-reducing terminus. These degrees of freedom give rhizobia the capacity to produce a multitude of Nod-factor variants (Miller and Oldroyd 2011). For instance, if each of 7 positions on the LCO backbone can have one of three possible decorations (including no decoration), there are 3^7^ = 2,187 possible variants, not counting length variation in the LCO backbone and acyl chains. The genetic architectures of Nod factors and Nod-factor-receptor genes both appear to favour rapid diversification. Known Nod-factor receptor genes are found in tandem arrays, and this might allow rapid evolution of these genes via recombination (Parniske and Downie 2003). Similarly, the diversity of *nod* genes allows rhizobia to add multiple and different decorations to the LCO backbone and to simultaneously produce multiple different Nod factor molecules (Miller and Oldroyd 2011).

However, what are the conditions under which passwords evolve via natural selection? Let us say that a bacterial lineage has co-evolved with a plant and that during this time, the symbiosis has progressed from an ancestral state of close association at the root surface (e.g. the plant root secretes carbohydrate compounds into its immediate surroundings and captures ammonium by diffusion and/or active transport) to the more intimate state of endosymbiosis, via the formation of an infection thread and root nodule. The benefit of evolving endosymbiosis is increased efficiency of nitrogen fixation in an oxygen-free environment, and the exclusion of free-riders is a possible additional benefit. *Nod*-gene duplication and mutation allowed rhizobia to evolve Nod factors of greater and greater complexity (more decorations added to the LCO chain). Nod factor does not have to be costly, except in the trivial sense that it needs some energy to be synthesized, and indeed Nod factor does not seem to have a *strategic* cost. However, Nod factor does need to be uniquely recognizable, which could explain why it is complex.

During the evolution of Nod factor, we assume that the host plant also had ancestral physiological mechanisms for shedding or withdrawing resources from worthless roots that have failed to take up fixed nitrogen (Partner Fidelity Feedback PFF, Weyl et al. 2010) or had even evolved punishment (Host Sanctions HS, Kiers et al. 2003). Either way, *rhizobial lineages with the correct Nod factor have been subjected to selection for nitrogen fixation and against pathogenicity*. The usefulness of passwords is that they can truthfully signal that a bacterial lineage shares a co-evolutionary history with a host lineage, and if that history is mutualistic, then the plant is selected to engage in symbiosis with bearers of the password. Thus, passwords signal a particular evolutionary history (an ‘identity’), and since that history is written into the bacterial symbiont’s genome, the genome enforces a particular behaviour.

This is what a person does with a password on a bank website: he credibly signals to the bank that they share a specific history of transactions, and if those transactions have been acceptable to the bank, there is a good chance that the person will continue to act acceptably, so the bank should allow continued transactions. (Passwords are an alternative to repeated games. By definition, players of repeated games build up interaction histories with other players and thus need to recognize individuals in order to apply the correct history to each player, but with passwords, it is possible for a member of a host lineage to recognize a member of a specific symbiont lineage, even if that pair of individuals has never met.)

For a password to be a reliable signal of identity, the password needs to be complex. Otherwise, a password that identifies one lineage could easily be evolved *de novo* by other lineages. Thus, our challenge becomes one of explaining why there is directional selection for signal complexity. We also need to explain ‘strictness’ in receivers, where strict means that the receiver rejects passwords with (too many) errors. It is the combination of a sufficiently complex signal and a sufficiently strict receiver that renders it essentially impossible for parasites to evolve a working password *de novo*.

### Passwords can evolve under soft selection

In the context of mutualistic interactions, the process we envisage leading to the evolution of cost-free but still honest signalling is the following. Consider a population of hosts harbouring only mutualistic symbionts. *Mutations in the symbionts* lead to a slightly different (simpler or more complex) password or to a parasitic phenotype. We assume that double mutations are negligible, meaning that a symbiont will not simultaneously evolve a new password and a newly parasitic phenotype. We also let *mutations in the hosts* lead them to accept a slightly different (simpler or more complex) password. Hosts accepting the original password will therefore accept both mutualistic and the new mutant parasitic partners, but hosts with mutations that cause them to accept a different password will reject parasitic symbionts, which still use the original password under the assumption that double mutations are negligible.

Mutant *hosts* will have higher fitness than the resident hosts if the benefit of avoiding parasites is higher than the cost of the difficulty of finding the rarer partners carrying the right mutant password (i.e. some mutant hosts fail to find a symbiont partner). If this is the case, mutant *symbionts* with the slightly different (simpler or more complex) passwords will also have higher fitness, because mutualistic symbionts with the original password suffer some of the cost of PFF or HS triggered by the co-colonising parasitic mutants (because the host expends energy to trigger PFF or HS and because of the opportunity cost of mutualists having lost out on colonisation opportunities to parasites). Then, we must only explain why the mutant symbionts with the more complex password have higher fitness than the mutants with the simpler password. We will show that, *provided that the hosts undergo soft selection*, the hosts that mutated to accept the more complex passwords have a selective advantage, and as a result, the signallers that mutated to slightly more complex passwords will also have a selective advantage. Repeating this scenario over time leads to increasingly complex passwords that are honest and cost-free.

The critical, non-intuitive step in this scenario is the contribution of *soft selection* (Buchholz 1922; Wallace 1981; Klekowski 1988) (**Figure 2**). Soft selection occurs when juveniles are produced in excess of available carrying capacity for adults. For instance, a plant produces many more seeds than there are patches in the environment to support adults, and as a result, juvenile populations are unavoidably bottlenecked by this exogenous mechanism, or to put it another way, when a parent loses some fecundity, its fitness remains virtually unaffected, since some offspring would have died no matter what. Since all offspring contain *de novo* mutations, which, if non-silent, are more likely to be deleterious than beneficial, soft-selected lineages evolve genes that are hypersensitive to mutation (i.e. antirobust); that is, soft-selected genes evolve to suffer large losses in function after mutation, causing the offspring that carry more mutations to be much more likely to die, which reduces competition with their more fortunate siblings that carry fewer mutations. Parental fitness benefits more from this filtering out of mutated offspring than is lost to reduced fecundity, since most offspring are destined to die anyway in a soft-selection scenario.

**Figure 2.**
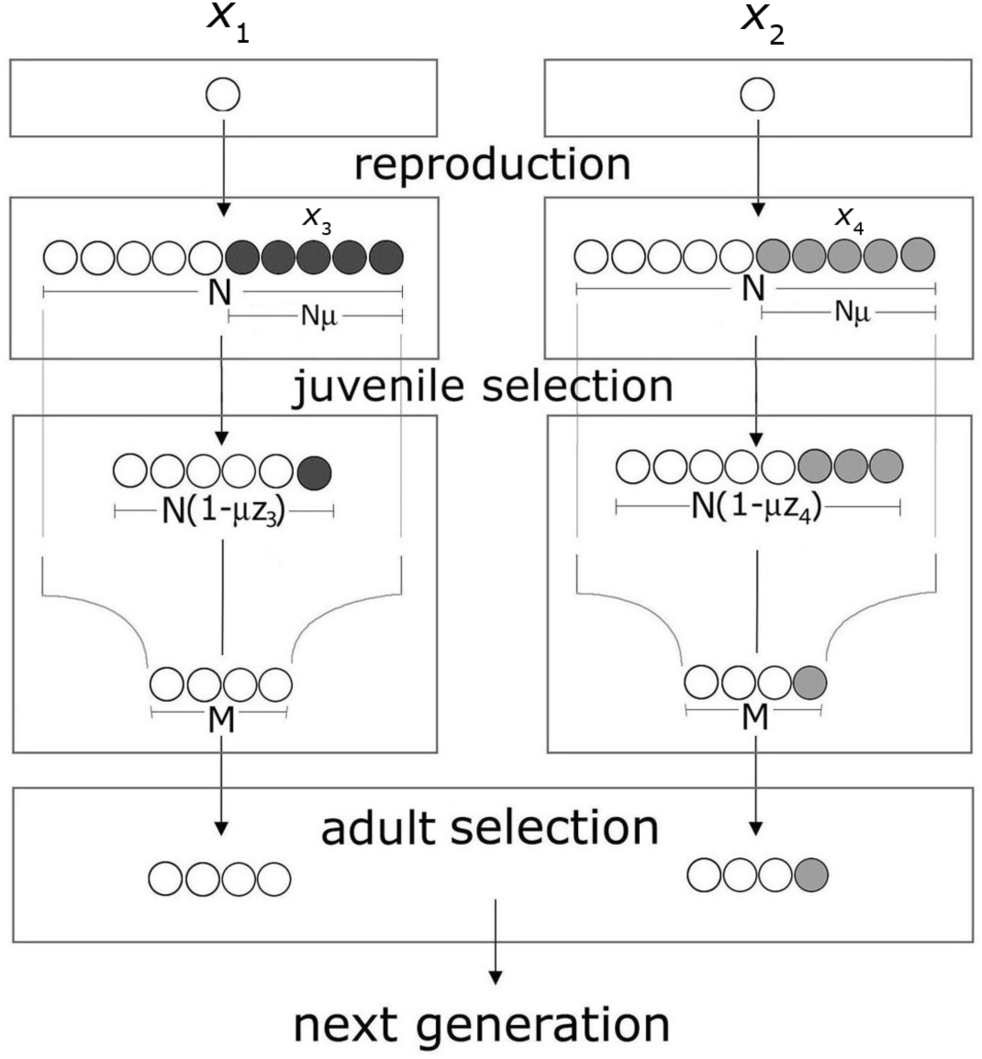
Soft selection leads to antirobustness. Alleles *x*_1_ and *x*_2_ have the same fitness. However, *x*_1_’s mutants *x*_3_ happen to have lower fitness than do *x*_2_’s mutants *x*_4_ (*z*_3_>*z*_4_, so (1-*μz*_3_) < (1-*μz*_4_)), and hence, *x*_1_ is less robust than is *x*_2_. If there is a juvenile selection stage, *x*_3_ mutants are more likely to be eliminated before the adult stage; if juvenile selection does not cause a loss of fecundity (because only a small fraction of the progeny goes on to the adult stage anyway), the adult progeny of *x*_1_ (M) will have a selective advantage over the adult progeny of *x*_2_, and *x*_1_ will increase in frequency.

In what follows, we use a population-genetics model to show that soft selection leads to the evolution of more complex passwords. We stick with the scenario in which the signaller is a bacterium and the receiver is a host plant, but the model applies in general to partner choice.

## Model

Consider a population of nitrogen-fixing rhizobial bacteria producing a password composed of separate elements *a, b, c*, etc. and allow each of these elements to arise or disappear in a stepwise fashion by a mutation that can add or remove one element (a mutation is denoted by adding or removing a letter from the password). For simplicity of notation, the order of the elements does not matter; that is, passwords of increasing complexity are denoted by *a, ab, abc*,…, etc., and *ab*=*ba, abc=acb*, etc. Note also that any element of a password can mutate (e.g. *abc* → *tbc* → *tbu*), but we ignore these scenarios here because our focus is on the origin of complexity *per se*, which we are representing by a longer password. In other words, we focus on explaining how to get from passwords of low to high complexity (*a* → *abcde*), and we ignore the diversification of passwords (*abcde* → *uewix*). Our way of denoting more complex passwords mimics the idea that Nod factor has evolved complexity by the proliferation of additional *nod* genes that have added more “decorations” to the basic LCO backbone (Miller and Oldroyd 2011).

In each generation, a fraction *μ*_P_ of the population produces a simpler password, and a fraction *μ*_P_ produces a more complex password. For instance, in a population of bacteria sending password *ab*, some mutants will send the password *a*, and some will send the password *abc*. Bacteria can also evolve to be parasitic by failing to export fixed nitrogen to the host plant. Hence, the population will also have a fraction *μ*_D_ of parasitic *ab*\* bacteria. Assuming that double mutations are rare, we ignore *a*\* and *abc*\* mutants.

Finally, we denote plants that accept a given password by capital letters; each plant type only accepts its own corresponding bacterium type (*AB* accepts only *ab* and *ab*\*; *A* accepts only *a*; *ABC* accepts only *abc*). This assumption of strictness in the plants is here made only for explanatory convenience, and we explicitly model the evolution of strictness in the next section. We start with a population fixed on *AB* plants and *ab* bacteria, with a low frequency of mutants *ABC, A, abc, a*, and *ab*\*.

### Signaller (Bacteria)

Selection coefficients *s*_**i**_ against the different bacterial types *i* in the **adult phase** are as follows (See **Appendix**).

*s*_ab_>0 because *AB* plants can be colonized by *ab* and *ab*\* bacteria. *ab* bacteria suffer some fitness loss due to *ab*\*’s cheating, as a consequence of the costs of host response against *ab*\* (direct costs of nodule senescence via HS or PFF in mixed nodules, indirect costs of senescence due to plant expenditure of energy, and opportunity costs of the plant failing to have hosted *ab* bacteria in all nodules, which were excluded via competition with *ab*\* bacteria for limited colonization opportunities).

*s*_ab*_>0 for the same reasons as above. If the plant reacts indiscriminately against *ab* or *ab*\*, then *s*_ab*_<*s*_ab_, since *ab*\* do not pay the cost of nitrogen fixation. If HS or PFF is preferentially targeted toward *ab*\*-inhabited nodules, then *s*_ab*_>*s*_ab_.

*s*_abc_=0 because there is no costly host response by *ABC* plants, since there are no *abc*\* bacteria. Strictly, *s*_abc_ = 0 when *abc* and *ABC* have zero additional difficulty in finding each other, relative to *ab* and *AB* finding each other. The assumption is reasonably upheld under soft selection because *ABC* plants make many *ABC* seedlings, which sample the bacterial population, and only some need to pick up *abc* bacteria. Note that this is an origin scenario for *abc* passwords, so a successful colonization of *ABC* plants only has to happen once in history for *abc* bacteria to start to be selected.

*s*_a_=0 for the same reason that *s*_abc_=0. There is no costly host response by A plants since there are no *a*\* bacteria in the population, due to our assumption of no double mutations.

In the simplest scenario, if we assume that there is no juvenile selection during the dispersal stage and that the (rare) mutant *a* and *abc* symbionts find their partners with no loss of fecundity, it is trivial to show that simple and complex passwords (*a* and *abc*) have the same fitness, which is greater than that of *ab* and *ab*\*, and that *a* and *abc* will have the same frequencies (0.5) at equilibrium, driving *ab* and *ab\** to extinction.

Of course, in a general scenario, there can be selection during dispersal, and rare types will be less likely to find the appropriate host, resulting in loss of fecundity. The juvenile selection coefficients *z*_*i*_ against the different bacterial types *i* measure the degree of selection against each allele i in the dispersal phase (during which the symbionts must find a suitable host).

The degree of soft selection *ϕ*_B_=*M*/*N* (where *N* is the number of offspring before juvenile selection and *M* is the maximum number of individuals that can go on to the adult phase) measures the buffering effect of soft selection. When *ϕ*_B_ =1, selection is hard, and losses during the dispersal phase result in reduced fitness. At the other extreme, when *ϕ*_B_=0, selection is soft, and any loss during the dispersal phase is fully recouped, resulting in no loss of fecundity.

To account for difficulty of finding rare hosts, we assume that there is no juvenile selection against the common alleles *ab* and *ab*\* but that the alleles with new passwords *a* and *abc* are less likely to find the appropriate but rare host (*A* and *ABC*). Therefore, *z*_ab_= *z*_ab*_=0 and *z*_a_, *z*_abc_>0. However, once the *a* and *abc* types have found a host, there is no selection against them in the adult phase, whereas there is selection in the adult phase against *ab*\*, which also induces selection against *ab* (as we outline above: *ab\** elicits nodule senescence in plants and compete for nodules, both of which therefore reduce fitness for *ab* as well).

Figure 3 shows the parameter space in which selection in the juvenile phase makes complex passwords increase in frequency. Complex passwords (*abc*) evolve as long as selection against the rare type during dispersal is lower (approximately) than selection against cheating mutants *ab*\* (and especially if selection occurs against *ab* as well) in the adult phase. Note that the degree of soft selection *ϕ*_B_ is largely irrelevant (left vs. right columns, Figure 3); that is, soft selection is not necessary for the complex password to evolve. Increasing *μ*_D_ and reducing *μ*_P_ increases the frequency of the *abc* allele. However, lower *μ*_D_ and higher *μ*_P_ enable the *abc* allele to increase in frequency for higher values of *z*_abc_.

**Figure 3.**
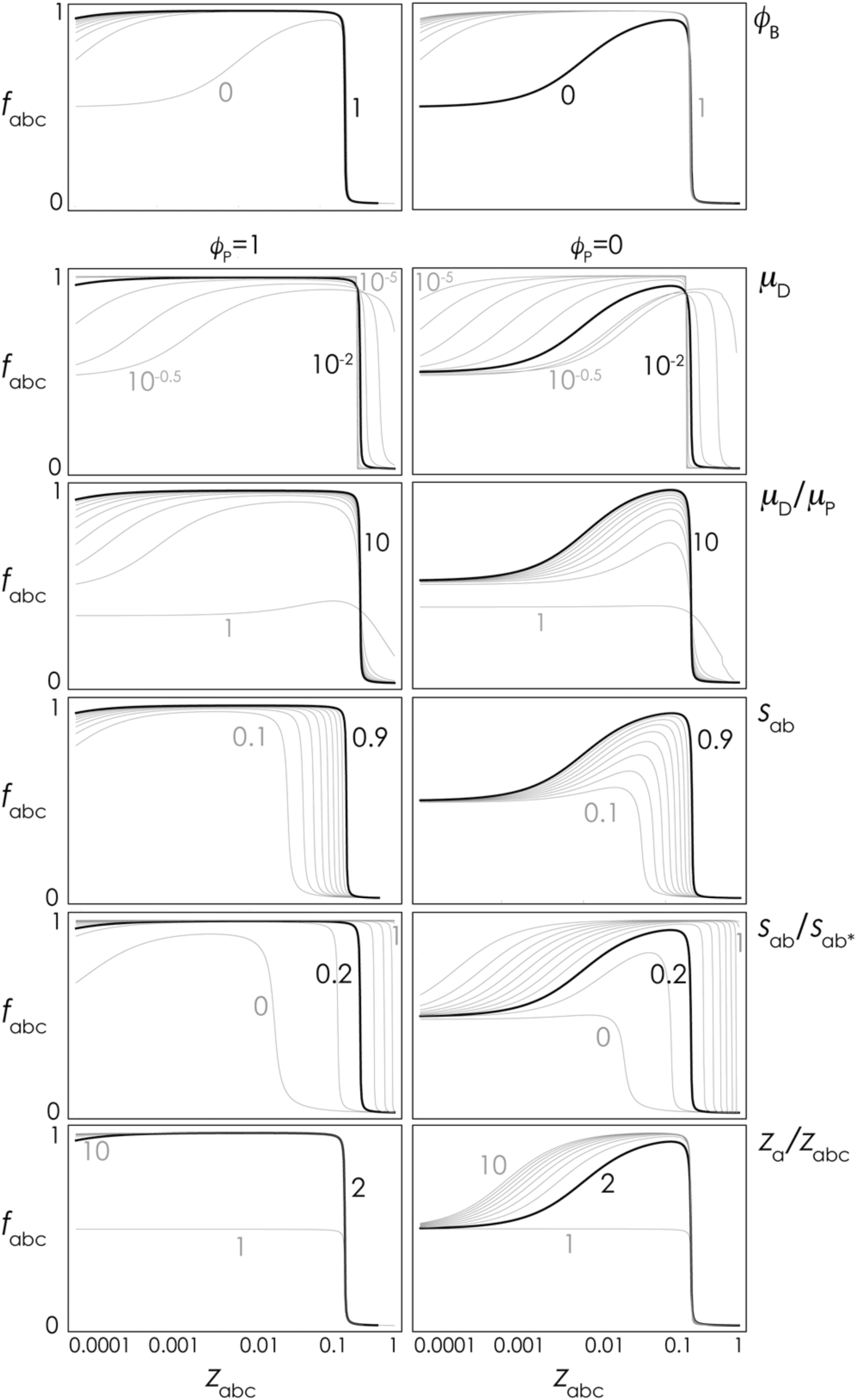
Frequency of the complex password (symbiont) at equilibrium. The equilibrium frequency of *abc* as a function of its coefficient of juvenile selection (*z*_*abc*_). The black curve shows the equilibria for *μ*_D_=10^-2^, *μ*_P_=10^-3^, *s*_ab*_=0.9, *s*_ab_=0.2*s*_ab*_, *z*_a_=2*z*_abc_ and for *ϕ*_B_=0 (right: no loss of fecundity in the juvenile phase) or *ϕ*_B_=1 (left: full loss of fecundity in the juvenile phase). The grey curves show the equilibria for other values of the parameters. There is no selection in the *adult* phase against *a* and *abc* (*s*_*a*_=*s*_abc_=0) and no selection in the *juvenile* phase against *ab* and *ab\** (*z*_ab_=*z*_ab*_=0).

*The complex password evolves even if there is mortality in the juvenile (dispersal) phase for the mutant passwords* a and abc; the *abc* allele increases in frequency (and may go to fixation) if *z*_a_>*z*_abc_. If *z*_a_=*z*_abc_, *a* and *abc* have frequency 0.5 at equilibrium (the exact *z*_a_/*z*_abc_ ratio is rather irrelevant, unless *ϕ*_B_=0).

In this scenario, both the simpler password *a* and the more complex password *abc* have an advantage against the original password because selection against *ab*\* and *ab* in the adult phase offsets the loss of fecundity in the juvenile phase (*z*_*i*_ >0, due to the difficulty in finding an appropriate host). Whether *a* or *abc* prevails is determined by whether the coefficients of juvenile selection for the two types are different, which depends on the rarity (hence on the fitness) of the respective hosts. If *ABC* plants increase in the population, then *abc* bacteria will have an advantage over *a* (*z*_a_>*z*_abc_), and selection will lead to more complex passwords. So now we have to explain why *ABC* plants evolve, rather than *A* plants.

In other words, so far, we have simply recovered the well-known finding that selection induced by parasites induces diversity in a population. We are left to explain why this diversity leads directionally to increased signal complexity. Our solution in the next section will invoke soft selection.

However, before going on, it is possible to outline a simpler but perhaps less general scenario for the evolution of password signalling. It might be the case that *s*_a_>0 (i.e. *a* suffers a fitness cost) if there are *a*\* bacteria in the population, which elicit PFF response in plants, and this response harms *a*, as in the argument for *ab* having *s*_ab_>0. This could occur if *a* passwords are simple enough that they can be evolved *de novo* by parasitic bacteria. In this case, it seems obvious that *abc* will go to evolutionary fixation, as long as the benefit of being a pathogen is not greater than the cost of PFF. Then, *ABC* plants will increase simply because they accept only *abc* bacteria, whereas *A* and *AB* plants accept some pathogenic bacteria (*a\** and *ab*\*).

In the following, let us retain the more conservative assumption that *s*_a_=0 (i.e. parasites induce diversity in the host, but this diversity is not biased towards higher or lower complexity). The next part of our explanation is to show why hosts evolving to accept higher complexity symbionts (*ABC*) have a selective advantage over hosts evolving to accept lower complexity symbionts (*A*).

### Receivers (Hostplants)

Let us consider a locus with four alleles: *ABC, A, ABC*^*^, *A*^*^, coding for the receiver’s recognition system. We only need to model two types (*A* and *ABC*) and their nonsense or missense mutants (*A*\*, *ABC*\*):

- Allele *ABC* always accepts password *abc*
- Allele *ABC*^*^ is a missense mutant of allele *ABC*
- Allele *A* always accepts password *a*
- Allele *A*^*^ is a missense mutant of allele *A*

The key assumption is that a mutant of *A* (*A*^*^) will be more likely to still accept its password *a* than a mutant of allele *ABC* (*ABC*^*^) will accept its password *abc*. In other words, we posit that allele A is *robust* to mutation and that allele *ABC* is *antirobust* to mutations. Note that robustness is unrelated to viability. *A* and *ABC* have exactly the same fitness (since *abc* and *a* bacteria have the same effect on the plant’s fitness); it is their mutants (*A*^*^ and *ABC*^*^) that have different fitnesses.

The mechanism behind our assumption could simply be that an *ABC* receiver system is necessarily made up of more numerous or complex molecules that interact with each other, since password *abc* is physically more complex than password *a*. Thus, even a mutation of ‘small effect’ in one component of *ABC* might prevent the *abc* signal molecule from fitting properly in the other component(s) of *ABC* and thus prevent the different receptor molecules from interacting correctly with each other to trigger a signalling cascade. A complexly interacting receptor is also more likely to be a strict receiver, since even a small change in the Nod factor would cause it fit differently in one or more of the receptor’s molecules and thus interfere with the interaction of those molecules.

We thus look for evidence that the Nod-factor receptor complex is composed of multiple molecules that need to interact with each other in order to trigger a proper signalling cascade. As it happens, there is considerable empirical support for this model (Oldroyd 2013). The Nod-factor receptor complex is made up of two separately produced receptor-like kinases (NFR1 and NFR5, using *Lotus japonicus* nomenclature), each of which carries extracellular LysM motifs that bind to the *N*-acetylglucosamine backbone of the Nod-factor backbone (Broghammer et al. 2012). The two receptor-like kinases heterodimerise *in vivo* (Madsen et al. 2011), and *mutation in either of the two kinases prevents rhizobial infection* (Radutoiu et al. 2003). Importantly, NFR5 is a non-functional kinase (this is known because it lacks essential protein subdomains), and thus, *NFR5 can act only via its interaction with NFR1* (Madsen et al. 2011). The activated NFR1/5 complex then appears to activate a third receptor-like kinase (SYMRK), which is necessary for downstream propagation of the infection pathway (Radutoiu et al. 2003). Finally, Morieri et al. (2013) have shown that the removal of just one acetyl group decoration from the *Sinorhizobium meliloti* Nod factor is enough to prevent calcium influx in its host, *Medicago truncatula*, and this failed influx prevents the initiation of an infection thread. Morieri *et al*. propose a model in which *only the correct Nod factor is able to bring about “cooperative interactions” between receptor proteins “such that the resulting interaction alters the kinase activity or specificity of the receptor complex*”, triggering the calcium influx that is needed for successful infection-thread initiation. In short, the Nod factor receptor is clearly a machine of many interdependent parts, and thus, of many points of failure.

The evidence that an *A* receptor would be more robust to mutation is sparser, because all known Nod-factor receptors (and Nod factors) are complex (Perret et al. 2000; Madsen et al. 2011; Miller and Oldroyd 2011; Broghammer et al. 2012). However, the Myc factors produced by arbuscular mycorrhizal (AM) fungi are structurally simpler (Maillet et al. 2011), and multiple, distantly related plant species will enter into symbiosis with the same AM fungal genotype, despite the reasonable expectation that the Myc receptor complexes from different plant lineages have mutated during plant diversification. However, because Myc-factor receptors are not yet characterised, it is not yet possible to rule out the alternative hypothesis that individual plant species produce multiple, species-specific Myc receptors.

Now, the evolution of password complexity requires that *ABC* increase in frequency over *A*. The question therefore is: why should *ABC* increase in frequency, given that alleles *ABC* and *A* are neutral? If anything, it seems that allele *A* should increase, since it is more robust to mutation and will therefore have a higher rate of back mutations from allele *A*^*^, whereas allele *ABC*^*^ is less likely to survive and will provide fewer back mutants to *ABC* (Wagner et al. 1997; Hermisson et al. 2002; de Visser et al. 2003).

We can see this effect in **Figure 4** left column, where we assume the standard hard-selection scenario in which juvenile mortality reduces adult fitness (*z*_*A*\*_=*z*_ABC*_=0). The frequency (*ƒ*_*A*_) of the robust allele *A* increases, provided that the selection coefficient s_ABC*_ against the defective mutant *ABC*^*^ is high enough (approximately higher than the mutation rate). If selection is too weak compared to the mutation rate, the differential amount of back mutations (to *ABC* and *A* from *AB*) is negligible, and the two alleles *A* and *ABC* maintain the same frequencies. If selection is strong enough, however, with no soft selection, the robust allele *A* increases in frequency over *ABC* because it receives more back mutations from *A*^*^ than *ABC* receives from *ABC*^*^.

**Figure 4.**
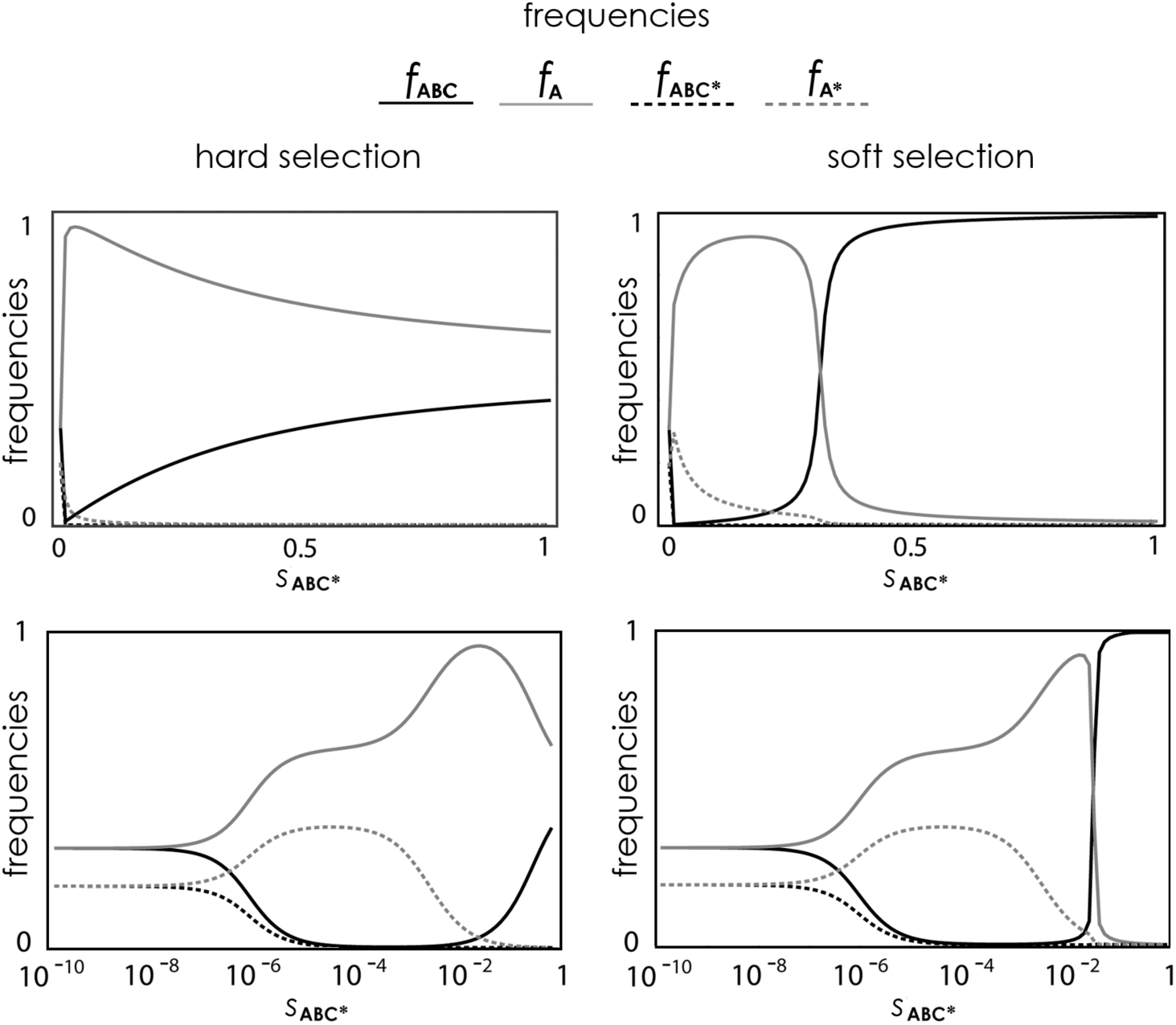
Frequency of the four types of receiver (host) at equilibrium. Equilibrium frequencies (*ƒ*_A_, *ƒ*_ABC_, *ƒ*_A*_, *ƒ*_ABC*_) of the four receptor alleles as a function of selection, with hard selection (left, *z*_i_=0) or with soft selection (right, *z*_i_=*s*_i_); *ϕ*_P_=1, *s*_A*_/*s*_ABC*_=1/10; *s*_A_=*s*_ABC_=0; *μ*_S_=10^-4^, *μ*_P_=10^-7^; *s*_A*_ and *s*_ABC*_ are selection coefficients against dysfunctional *A* and *ABC* alleles, which arise with probability *μ*_S_. The horizontal axis in the two bottom panels is logarithmic, to highlight the equilibrium frequencies with weak selection.

However, *with soft selection*, antirobust alleles increase in frequency over robust alleles (**Figures 4**, **5**, **Appendix 1**). In fact, if the selection coefficients are high enough, the antirobust allele *ABC* can go to fixation. Strong selection is necessary, but not high mutation rates. Contrary to the evolution of robustness, the magnitude of the mutation rate does not make any relevant difference to the evolution of anti-robustness because the driving force is not the rate of back mutation, but the soft-selection process that eliminates the non-functional *ABC* mutants (Otto and Hastings 1998). Lower rates of mutation to non-functional *ABC* mutants do enable complex passwords to evolve for lower levels of soft selection (**Figure 5**).

**Figure 5.**
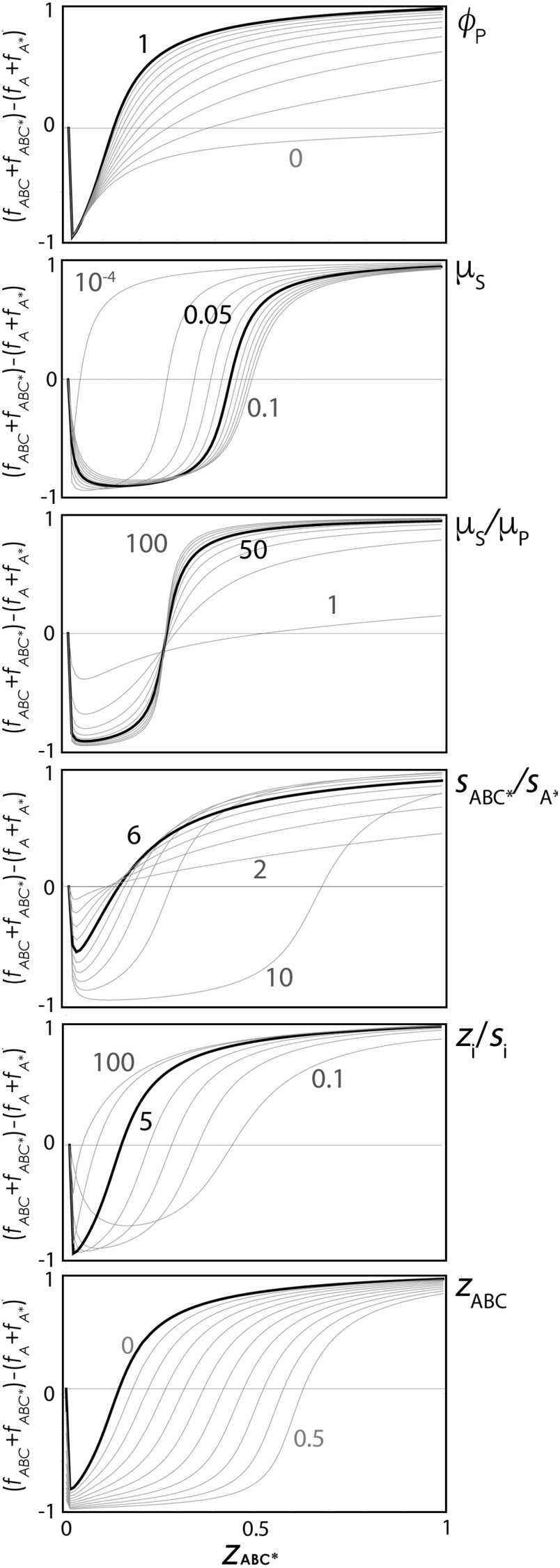
Frequency of the antirobust recognition system (host) at equilibrium. Equilibrium frequency of *ABC* as a function of its coefficient of juvenile selection (*z*_*ABC*_). The black curve shows the equilibria for parameter values *ϕ*_P_=0, *s*_ABC*_/*s*_A*_=10, *s*_i_/*z*_i_ =1, *μ*_S_=10^-2^, *μ*_S_/*μ*_P_=50. The grey curves show the equilibria for other values of the parameters. There is no selection in the adult phase against *A* and *ABC* (*s*_A_=*s*_ABC_=0) and no selection in the juvenile phase against *A* (*z*_A_= 0).

The fraction of complex passwords at equilibrium increases as the ratio between the mutation rates for passwords and the mutation rate to defective alleles decreases (**Figure 5**), because if the mutation rate to defective alleles is too high, the advantage derived from soft selection is offset by partial loss of fecundity. A similar effect is observed for *z*_*i*_/*s*_*i*_ (**Figure 5**), because when juvenile selection is stronger, defective alleles are more easily eliminated from the offspring (hence, the original antirobust allele is more likely to increase in frequency). The influence of *s*_*ABC*\*_/*s*_*A*\*_ is more complex, as it depends on the value of *z*_*ABC*\*_ (**Figure 5**). Note that soft selection can lead to an increase in the complex (antirobust) allele even if complexity itself has a cost (*z*_*ABC*_ >0).

## Discussion

### Passwords: an honest, verifiable-information signal

Game theory recognizes three classes of honest signalling models: costly signals, cheap talk, and verifiable information. Passwords can be thought of as a variant of the verifiable-information class. By virtue of their complexity and thus the low likelihood that they can evolve *de novo*, they serve as signals of lineage identity that are self-evidently true, so long as the receiver can recognise the password. In the context of horizontally transmitted mutualisms, passwords can evolve to reliably signal a shared coevolutionary history, and a coevolutionary history between mutualistic lineages strongly implies that the individual sending the password is itself a mutualist or a recent descendent of a mutualist.

We emphasise that the role of signalling is only to allow the receiver to distinguish different types, here, mutualistic and parasitic so that the receiver can associate preferentially with one of them. The source of the natural selection for mutualistic types in the first place is some form of partner fidelity feedback or host sanctions, and the source of parasitic types is mutation. There is a superficial similarity of password signalling with green-beard signals (Jansen and van Baalen 2006), which allow kin to identify each other, but green beards are a within-species mechanism, and passwords can act between species. Green-beard signals can also be simple, due to their linkage with cooperation loci.

In short, we argue that password signalling allows hosts to engage in successful Partner Choice. As it happens, it has been shown experimentally that legume plants are able to associate preferentially with ‘more mutualistic’ (nitrogen-fixing) rhizobial bacteria (Heath and Tiffin 2009; Gubry-Rangin et al. 2010; Sachs et al. 2010). Importantly, these studies used mutualistic and parasitic rhizobial bacteria that had been isolated from the same soil as the host plant, and we predict that the rejected parasitic bacterial lineages used in these experiments were producing Nod factors that had diverged from the mutualistic lineages that were accepted.

Our proposed scenario for the evolution of password signalling derives from the observation that plants are subject to a non-trivial degree of soft selection, since plants generally make many more juveniles than can possibly grow into adults. As a result, juvenile mortality in plants, due to some combination of failure to find bacterial symbionts and of selection against missense mutations, does not result in fitness loss.

We then posit that mutations in the genes for complex Nod-factor receptors (*ABC*) are inherently more likely to result in non-functional receptors than are mutations in the genes for simple-signal receptors (*A*), because more complex Nod-factor receptors are likely constructed from more interdependent parts. Thus, only fully functional *ABC*-receptors are likely to be represented in adult plants, because *ABC*-mutant juveniles will have died due to an inability to recruit rhizobia. In contrast, when the genes for *A*-type receptors mutate, the receptors are more likely to retain some function because they are simple, and some of these lower fitness mutants will thus be represented in the adult stage. Competition between *ABC* and *A* adults will then favour *ABC*, and thus, complex rhizobial passwords (*abc*) will also be favoured, and the system will evolve toward complex signals of identity. After enough rounds, Nod factor will have evolved to a high enough degree of complexity that it will be essentially impossible for a bacterium to evolve a working Nod factor *de novo*.

We also recall our first, and simpler, scenario for the evolution of complex signalling passwords, which relies on the possibility that simple Nod factors can evolve *de novo* in non-mutualistic bacteria (a^*^). In this situation, bacteria that evolve more complex passwords (*abc*) are favoured over those that evolve simpler passwords (*a*), as the former will find themselves in parasite-free hostplants, at least until *abc* bacteria evolve parasitic behaviour *abc*\*. This scenario also relies on soft selection, in that many *ABC* juvenile plants will die before finding a suitable *abc* partner, but as long as there are lots of *ABC* juveniles, some will be successful, and these will form the next generation.

### Limits to complexity in passwords

In either scenario, we expect a natural upper limit to the complexity of passwords because there will be physical limits on the reliable functioning of complex-signal receptors (*s*_*ABC*_ > 0), and receptors that evolve beyond these limits will likely fail to perceive any symbionts, which reduces the effect of soft selection (**Figure 5**). Thus, the evolution of complexity in passwords cannot escape indefinitely from the evolution of parasitic genotypes within rhizobial lineages. There must also be mechanisms to senesce nodules that have been colonised by parasites (Kiers et al. 2003; Weyl et al. 2010). And of course, such mechanisms were necessary to proliferate the mutualistic genotypes of rhizobia in the first place, or there would have been no mutualistic lineages for the plant to recognise.

Once a combination of password signalling and selective nodule senescence has evolved, mutualistic strains of rhizobia should grow to dominate soils. As a result, it is possible to imagine situations where some hosts will evolve to relax the strictness of association or evolve to accept multiple passwords. As one example, some leguminous tree species, and the non-legume Cannabaceae plant genus *Parasponia*, are early-successional species that colonise low-nutrient soils, and they can be colonised by multiple rhizobial genera, including strains from different continents (Behm et al. 2014). Under such conditions, we expect that the risk of being colonised by non-productive or even pathogenic bacteria is outweighed by the benefit being able to fix nitrogen. It will be interesting to see if these species have evolved multiple Nod factor receptors, or if their receptors are less strict (which should make them more robust to mutation). Indeed, in *Parasponia andersonii*, it appears that the latter might be true, because this species uses the same receptor for both Myc and Nod factors (Op den Camp et al. 2011).

Myc factors, which consist of simple, almost entirely undecorated LCOs (Maillet et al. 2011), provide an interesting counterexample to complex Nod factors. Why have AM fungi not evolved complex passwords? Part of the answer is likely due to the fact that plants are colonized by multiple AM fungal species, and by doing so, plants make the fungi compete for plant carbon, thereby reducing the carbon cost of AM-provided phosphorous (Argüello et al. 2016). A plant that evolved a more complex receptor would reduce its diversity of fungal partners and thus reduce the number of competing fungal suppliers. It is also possible that each AM fungal genotype benefits from colonizing multiple plant species, if plants vary temporally in the photosynthate that they are able to transfer to their fungal partners. An individual AM fungus that evolved a more complex Myc factor recognised only by the rare plant genotype that had also evolved a matching receptor would not be able to create to create such networks.

### Extensions to the model

Our model has some limitations. First, it only shows that more complex passwords will, under certain conditions, increase in frequency over less complex passwords, without explicitly showing that they will become more and more complex over time. Increasing complexity, however, is obvious if we assume that more complex alleles arise by mutation after the fixation of the complex password (i.e. *ABC* will increase in frequency over *AB*, then *ABCD* will increase in frequency over *ABC*, and so on). If mutants arise before fixation, the dynamics will be more complex, but the logic remains the same: more complex passwords will have an advantage over less complex ones under soft selection. Second, our model does not allow for the fitness of an allele being dependent on the proportions of its partner in the population. It does, however, allow juvenile selection against rare alleles (the phase in which partner choice occurs). While it might be interesting to analyse frequency-dependent, soft selection coefficients, there seems to be no compelling reason why this should select against signal complexity. Third, we have ignored the fact that, as alleles become more complex, asymmetries in mutation rates may arise (e.g. mutations from *ABC* to *A* might be more likely than mutations from *ABC* from *A*). Preliminary results with asymmetric mutation rates, however, did not reveal significant differences in the results, as long as the differences are not extreme.

### Password signalling in mate recognition and defensive-symbiont acquisition

Soft selection occurs in practically all vascular plants, and also in many cryptogamic plants and in animals (Buchholz 1922; Wallace 1981; Klekowski 1988; Archetti 2009). Although detailed treatments are outside the scope of this paper, we hypothesise that password signalling can evolve in other recognition systems. For example, polymorphic toxin systems (PTS) comprise complex, multi-domain molecules that exhibit high levels of allelic diversity. Hillman and Goodrich-Blair (2016) have proposed that eukaryotic hosts can directly identify suitable defensive symbionts to acquire by sensing the PTS produced by those symbionts, and they review evidence that hosts produce PTS-receptors that are specific to particular symbiont lineages. Another possible class of password signals are post-mating recognition systems in gametes, which could require either a complex or a simple signal to differentiate conspecifics from heterospecifics. If the complex recognition system is more antirobust to mutation, then gametes that have suffered mutation will die unmated. However, those that survive will only have accepted conspecifics. In contrast, gametes that accept a simple signal might be robust to mutation and thus accept heterospecifics, producing hybrids. Under the twin assumptions that soft selection is acting (most juveniles die before achieving adulthood) and that hybrids have lower fitness, there could be selection for a mate-recognition system that requires a complex signal.

## Acknowledgments

We thank István Scheuring and Gergely Boza for illuminating discussions. DWY was supported by the National Natural Science Foundation of China (41661144002), the Ministry of Science and Technology of China (2012FY110800), the University of East Anglia, and the State Key Laboratory of Genetic Resources and Evolution at the Kunming Institute of Zoology (GREKF13-13, GREKF14-13, and GREKF16-09). JBM is supported by a fellowship from the University of East Anglia and the Eastern Academic Research Consortium.

## Appendix

## Plants

Consider a locus with four alleles: *ABC, A, ABC*^*^, *A*^*^ (with frequencies *ƒ*_ABC_, *ƒ*_A_, *ƒ*_ABC*_, *ƒ*_A*_, respectively:

We assume that alleles *A* and *ABC* can mutate to each other with the same probability μ_P_ and that each can mutate to dysfunctional alleles (respectively *A*\* and *ABC*\*) with probability μ_S_. We assume that all alleles have the same total mutation rate, hence *A*\* and *ABC*\* have other mutants (*A*^0^ and *ABC*^0^) with frequency *μ*_P_ that have zero fitness -- see **Table A1**.

**Table A1.** Offspring and fitness for the four plant types.

|  | fitness | offspring |  |  |  |
| --- | --- | --- | --- | --- | --- |
| | | $f_{ABC}$ | $f_A$ | $f_{ABC^*}$ | $f_{A^*}$ |
| $f_{ABC}$ | $1-s_{ABC}$ | $(1-\mu_S-\mu_P)(1-z_{ABC})/\alpha_P$ | $\mu_P(1-z_A)/\alpha_P$ | $\mu_S(1-z_{ABC^*})/\alpha_P$ | 0 |
| $f_A$ | $1-s_A$ | $\mu_P(1-z_{ABC})/\beta_P$ | $(1-\mu_S-\mu_P)(1-z_A)/\beta_P$ | 0 | $\mu_S(1-z_{A^*})/\beta_P$ |
| $f_{ABC^*}$ | $1-s_{ABC^*}$ | $\mu_S(1-z_{ABC})/\chi_P$ | 0 | $(1-\mu_S-\mu_P)(1-z_{ABC^*})/\chi_P$ | 0 |
| $f_{A^*}$ | $1-s_{A^*}$ | 0 | $\mu_S(1-z_A)/\delta_P$ | 0 | $(1-\mu_S-\mu_P)(1-z_{A^*})/\delta_P$ |

The recurrence equations for this system are:

T·*ƒ*_ABC_’ = *ƒ*_ABC_(1-s_ABC_)(1-*μ*_S_-*μ*_P_)(1-z_ABC_)/α_P_+*ƒ*_A_(1-s_A_)*μ*_P_(1-z_ABC_)/β _P_+*ƒ*_ABC*_(1-s_ABC*_)*μ*_S_(1-z_ABC_)/*χ* _P_

T·*ƒ*_A_’ = *ƒ*_ABC_(1-s_ABC_) *μ*_P_(1-z_A_)/α _P_+*ƒ*_A_(1-s_A_)(1-*μ*_S_-*μ*_P_)(1-z_A_)/β _P_+*ƒ*_A*_(1-s_A*_)*μ*_S_(1-z_A_)/δ_P_

T·*ƒ*_ABC*_’ = *ƒ*_ABC_(1-s_ABC_) *μ*_S_(1-z_ABC*_)/α_P_+*ƒ*_ABC*_(1-s_ABC*_)(1-*μ*_S_-*μ*_P_)(1-z_ABC*_)/*χ* _P_

T·*ƒ*_A*_’ = *ƒ*_A_(1-s_A_) *μ*_S_(1-z_A*_)/β _P_+*ƒ*_A*_(1-s_A*_)(1-*μ*_S_-*μ*_P_)(1-z_A*_)/δ _P_

where T is a normalizing factor obtained by summing the right-hand side of the four above equations; α_P_, β_P_, χ_P_, δ_P_ are normalizing factors of the offspring frequencies (see Table 1) ; ϕ_P_=*M*/*N* is the degree of soft selection, where *N* is the number of offspring before soft selection (the same for all alleles) and *M* (<*N*) is the maximum number of individuals that can go on to the adult phase after soft selection. *N*_i_’ is the number of offspring of an individual with allele *i* after soft selection:

*N*_ABC_’ = *N*[(1-*μ*_S_-*μ*_P_)(1-*z*_ABC_)+ *μ*_P_(1-*z*_A_)+ *μ*_S_(1-*z*_ABC*_)]

*N*_A_’ = *N*[(1-*μ*_S_-*μ*_P_)(1-*z*_A_)+*μ*_P_(1-*z*_ABC_)+*μ*_S_(1-*z*_A*_)]

*N*_ABC*_’ = *N*[(1-*μ*_S_-*μ*_P_)(1-*z*_ABC*_)+*μ*_S_(1-*z*_ABC_)]

*N*_A*_’ = *N*[(1-*μ*_S_-*μ*_P_)(1-*z*_A*_)+*μ*_S_(1-*z*_A_)]

The effect of soft selection on offspring frequencies is that frequencies are normalized (because we are assuming no loss of viability) after juvenile selection by dividing them by the total frequencies of the surviving offspring (α_P_, β_P_, *χ*_P_, δ_P_ as appropriate, see **Table 1**). Individuals with these normalized frequencies go on to the adult phase, where another round of (hard) selection occurs. Selection in the juvenile phase has no effect on fecundity if *z*_*i*_<1-ϕ.

The equilibrium frequencies of the four alleles can be found by specifying the parameters *μ*_S_, *μ*_P_, *s*_i_ and *z*_i_ for the system above and calculating the leading eigenvector. The situation in which allele 1 is less robust than allele 2, is given by *s*_ABC*_>*s*_A*_>0 and *z*_ABC*_>*z*_A*_>0.

## Bacteria

Consider a locus with four alleles: *a, ab, abc, ab*^*^, with frequencies *ƒ*_a_, *ƒ*_ab_, *ƒ*_abc_, *ƒ*_ab*_, respectively

- Allele *ab* codes for a password of intermediate complexity
- Allele *a* codes for a password of lower complexity
- Allele *abc* codes for a password of higher complexity
- Allele *ab*^*^ codes for a password of intermediate complexity in a cheater bacterium

We assume that alleles *a* and *abc* can mutate to *ab* only among the alleles that can enter plants but that the total mutation rate is the same for all alleles. Hence, *a* and *abc* also have other mutants with frequency *μ*_P_ that produce either a password that has no match in the plant population or a defective bacterium (and therefore zero fitness) – see **Table A2**.

**Table A2.** The offspring and fitness of the four types of bacteria

| freq. | fitness | offspring |  |  |  |
| --- | --- | --- | --- | --- | --- |
| | | $f_{ab}$ | $f_a$ | $f_{abc}$ | $f_{ab}^*$ |
| $f_{ab}$ | $1-s_{ab}$ | $(1-2\mu_P-\mu_D)(1-z_{ab})/\alpha_B$ | $\mu_P(1-z_a)/\alpha_B$ | $\mu_P(1-z_{abc})/\alpha_B$ | $\mu_D(1-z_{ab}^*)/\alpha_B$ |
| $f_a$ | $1-s_a$ | $\mu_P(1-z_{ab})/\beta_B$ | $(1-2\mu_P-\mu_D)(1-z_a)/\beta_B$ | 0 | — |
| $f_{abc}$ | $1-s_{abc}$ | $\mu_P(1-z_{ab})/\chi_B$ | 0 | $(1-2\mu_P-\mu_D)(1-z_{abc})/\chi_B$ | 0 |
| $f_{ab}^*$ | $1-s_{ab}^*$ | $\mu_D(1-z_{ab})/\delta_B$ | 0 | 0 | $(1-2\mu_P-\mu_D)(1-z_{ab}^*)/\delta_B$ |

The recurrence equations for this system are:

T·*ƒ*_ab_’=*ƒ*_ab_(1-*s*_ab_)(1−2*μ*_P_-*μ*_D_)(1-*z*_ab_)/α_B_+*ƒ*_a_(1-*s*_a_)*μ*_P_(1-*z*_ab_)/β_B_+*ƒ*_abc_(1-*s*_abc_)*μ*_P_(1-*z*_ab_)/*χ*_B_ + *ƒ*_ab*_(1-*s*_ab*_)*μ*_D_(1-*z*_ab_)/δ_B_

T·*ƒ*_a_’=*y*_ab_(1-*s*_ab_) *μ*_P_(1-*z*_a_)/α_B_+*ƒ*_a_(1-*s*_a_)(1−2*μ*_P_-*μ*_D_)(1-*z*_a_)/β_B_

T·*ƒ*_abc_’=*y*_ab_(1-*s*_ab_) *μ*_P_(1-*z*_abc_)/α_B_+*ƒ*_abc_(1-*s*_abc_)(1−2*μ*_P_-*μ*_D_)(1-*z*_abc_)/χ_B_

T·*ƒ*_ab*_’=*y*_ab_(1-*s*_ab_) *μ*_D_(1-*z*_ab*_)/α_B_+*ƒ*_ab*_(1-*s*_ab*_)(1−2*μ*_P_-*μ*_D_)(1-*z*_ab*_)/δ_B_

Where α_B_, β_B_, χ_B_, δ_B_ are normalizing factors of the offspring frequencies (see Table 2) and ϕ_B_ =*m*/*n* is the degree of soft selection, where *n* is the number of offspring before soft selection (the same for all alleles) and *m* (<*n*) is the maximum number of individuals that can go on to the adult phase after soft selection. *n*_*i*_’ is the number of offspring of an individual with allele *i* after soft selection:

*n*_ab_’ = *n*[(1−2*μ*_P_-*μ*_D_)(1-*z*_ab_)+*μ*_P_(1-*z*_a_)+*μ*_P_(1-*z*_abc_)+ *μ*_D_(1-*z*_ab*_)]

*n*_a_’ = *n*[(1−2*μ*_P_-*μ*_D_)(1-*z*_a_)+ *μ*_P_(1-*z*_ab_)]

*n*_abc_’ = *n*[(1−2*μ*_P_ -*μ*_D_)(1-*z*_abc_)+ *μ*_P_(1-*z*_ab_)]

*n*_ab*_’ = *n*[(1−2*μ*_P_-*μ*_D_)(1-*z*_ab*_)+ *μ*_D_(1-*z*_ab_)]

